# GraFT: Graph Filtered Temporal Dictionary Learning for Functional Neural Imaging

**DOI:** 10.1101/2021.05.24.445514

**Authors:** Adam S. Charles, Nathan Cermak, Rifqi Affan, Ben Scott, Jackie Schiller, Gal Mishne

**Affiliations:** Department of Biomedical Engineering, Kavli Neuroscience Discovery Institute, Center for Imaging Science, and Mathematical Institute for Data Science, Johns Hopkins University, Baltimore, MD 21287 USA; Technion – Israel Institute of Technology, Haifa, Israel, 31096; Department of Psychological and Brain Sciences, Boston University, MA 02215 USA; Halicioğlu Data Science Institute and the Neurosciences Graduate Program, UC San Diego, CA 92093 USA

**Keywords:** dictionary learning, sparse coding, calcium imaging, two-photon microscopy, re-weighted *ℓ*_1_

## Abstract

Optical imaging of calcium signals in the brain has enabled researchers to observe the activity of hundreds-to-thousands of individual neurons simultaneously. Current methods predominantly focus on matrix factorization and aim at detecting neurons in the imaged field-of-view, and then inferring the corresponding time-traces. The explicit locality constraints on the cell shapes additionally limits the applicability to optical imaging at different scales (i.e., dendritic or widefield data). Here we present a new method that frames the problem of isolating independent fluorescing components as a dictionary learning problem. Specifically, we focus on the time-traces, which are the main quantity used in scientific discovery, and learn the dictionary of time traces with the spatial maps acting as the presence coefficients encoding which pixels the time traces are active in. Furthermore, we present a novel graph filtering model which redefines connectivity between pixels in terms of their shared temporal activity, rather than spatial proximity. This model greatly eases the ability of our method to handle data with complex non-local spatial structure, such as dendritic imaging. We demonstrate important properties of our method, such as robustness to initialization, implicitly inferring number of neurons and simultaneously detecting different neuronal types, on both synthetic data and real data examples. Specifically, we demonstrate applications of our method to calcium imaging both at the dendritic, somatic, and widefield scales.

## I. Introduction

Functional optical imaging has become a vital technique for simultaneously recording large neural populations at single cell resolution hundreds of micrometers beneath the surface of the brain in awake behaving animals [1]–[3]. This class of techniques has become quite extensive, including one-two- and three-photon imaging [1], [4], [5], imaging at dendritic, somatic and widefield scales [1], [6]–[8], and imaging of various indicators of neural activity, including calcium, voltage etc. [9]–[11]. Of these methods, calcium imaging (CI) via two-photon microscopy has emerged as a dominant modality providing a practical method to optically record the calcium concentrations in neural tissue that is intrinsically representative of neural activity [1]–[3].

Given the large, and ever-growing, size of CI datasets, recent advances in automated CI analysis have sought to remove the need for traditional manual annotation of such data. To date, automatic analysis of CI primarily focused on factoring the recorded video into a two sets of variables: the spatial profiles^1^ representing the area that a neuron occupies, and the corresponding temporal fluorescence activity traces. Novel extensions, such as volumetric imaging [12], [13] and widefield imaging [8], [14], [15], promise to even further increase the dimensionality of such data and the spatiotemporal statistics that must be leveraged for accurate demixing.

As somatic imaging is the most prevalent, CI demixing algorithms have largely incorporated spatio-temporal regularization based on the statistics of somatic components to improve cell-finding. Specifically, somatic components are often well localized with sparse activity. Some methods, such as deep learning based methods [16]–[18], active contours [19], and spectral embeddings [20]–[22], include these statistics implicitly. Other methods, largely based on regularized non-negative matrix factorization, explicitly include this information in the optimization cost function [23]–[28]. These models are often prone to overfitting noise and motion artifacts, so the additional regularization terms capturing the spatial cohesion and sparse firing are necessary for interpretable results [23], [27]–[31].

More recently, variants of optical imaging have aimed to expand the scope of accessible brain signals by imaging both larger and smaller neural structures. At one end of this spectrum, zooming in enables the imaging of dendritic and spine structures, which captures how individual neurons communicate [32], [33]. Dendritic imaging, while also having sparse temporal statistics, can have long, thin spatial profiles that span the entire field-of-view (FOV). At the other end, cortex-wide (i.e., widefield) imaging can be achieved at resolutions too coarse to isolate activity signals of individual neurons, but instead can capture brain-wide activity patterns in freely moving animals [14]. At both scales, the spatial statistics of the sought-after components differ significantly from somatic imaging, and require new approaches that can be seamlessly applied across modalities.

Here we propose two major changes to improve the accuracy and scope of inferring fluorescing components from CI data. First, following our recent work [34], we refocus the goal of the problem onto finding the time-traces of each component.

Mathematically, we reorient the matrix factorization model formulation from

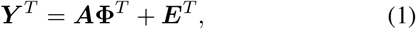

where 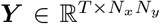, is the *N_x_* × *N_y_* pixel FOV sampled at *T* time-points, 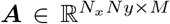 and **Φ** ∈ ℝ^*T*×*M*^ are the *M* spatial profiles and corresponding time-traces for each component, and 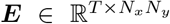 is *i.i.d.* Gaussian sensor noise^2^, to the model’s transpose

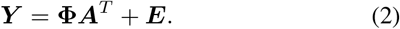

The goal for this step is not to change the model, as in fact both formulations are trivially identical, however we aim to change the philosophy with which calcium imaging is viewed, putting the emphasis on the time-traces which are the critical component to learning how neural activity is linked to behavior, stimuli, learning, etc. This philosophical shift strongly recommends an algorithmic shift to a dictionary learning paradigm, where we treat the time-traces as the dictionary. Furthermore, the spatial regularization becomes a natural extension, correlating the dictionary decomposition between pixels [35].

The second contribution is generalizing the rigid spatial grid of the FOV to a more flexible graph model over the pixels. Th additional graph modeling layer serves to remedy the fact that in many imaging scales, the typical assumptions of co-localized spatial profiles (i.e., neighboring pixels are likely composed of similar components) no longer holds, e.g., in dendritic imaging (Fig. 1A). The graph essentially redefines what “neighborhood” means, pulling together, or moving apart, pixels based on their temporal correlation structure and not their spatial adjacency. The graph-based regularization is used for deciding how the sparse coefficients (i.e., the spatial maps) are correlated as a function of the pixel-wise temporal correlations.

**Fig. 1.**
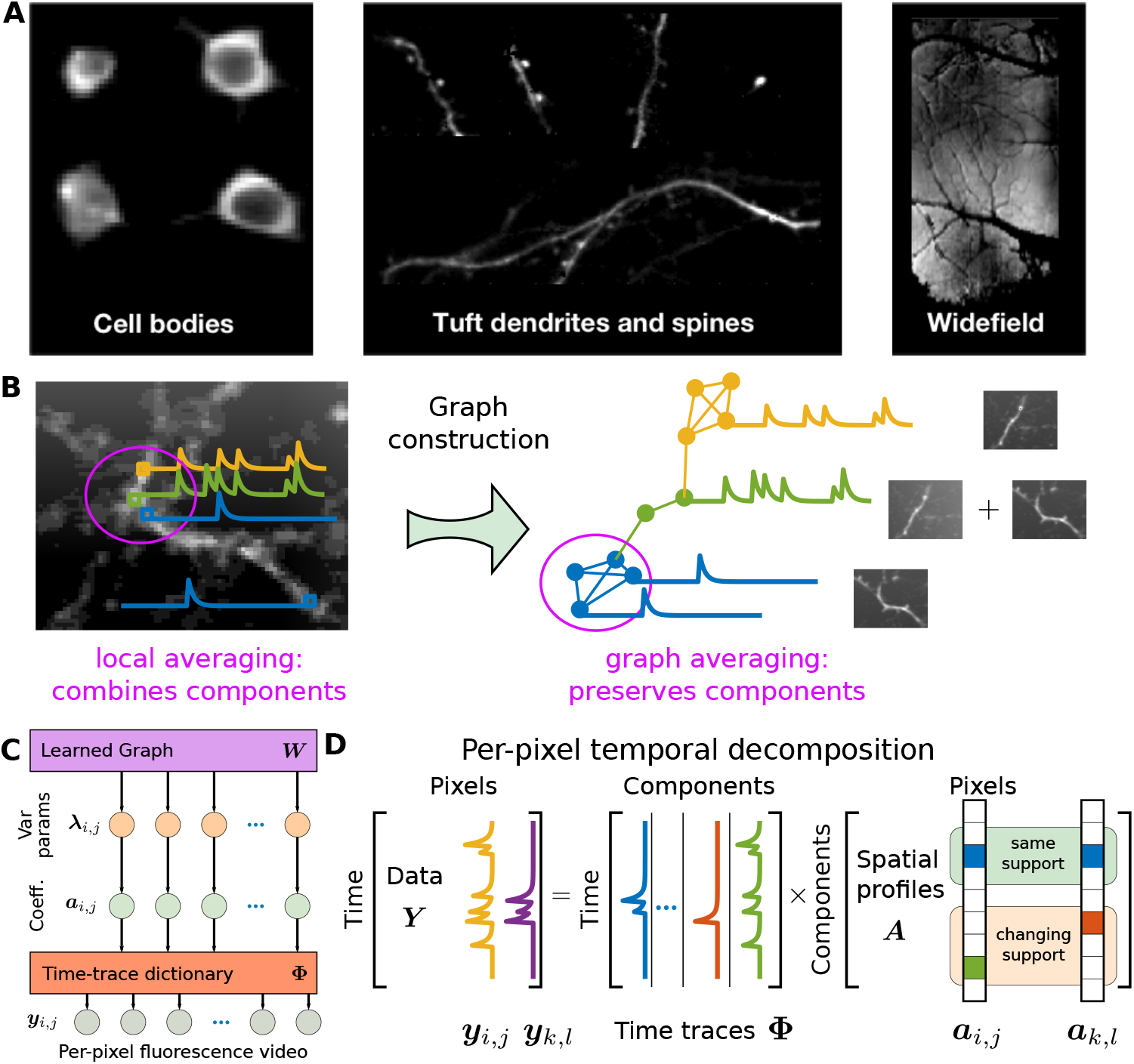
Basic concept of graph-based dictionary learning for calcium image analysis. A: Optical imaging of different neural components at different scales requires having flexible spatio-temporal regularization. B: In calcium imaging videos, distant pixels might be highly correlated while neighboring pixels might be very different. For example consider the depicted example of two crossing dendrites. Any technique based on local averaging to extract useful signals is thus prone to corruption by competing signals. By reorganizing the pixel relationships into a data-drive graph, local averaging (now defined by the graph) can make much more judicious use of temporal correlations between pixels and extract better signal estimates. C: The graphical model of GraFT dictionary learning organizes variables into three layers: a set of variance parameters interconnected by the data-driven graph ***W***, a set of sparse coefficients conditioned on the parameters, and the observed fluorescence video which is linearly generated by the coefficients through the dictionary **Φ**. D: As a matrix factorization, our model emphasizes the role of time-traces within the graph-based framework. The model thus focuses on pixel-wise decomposition into a dictionary of component time-traces.

Taken together, these two complementary changes give us a flexible new tool for automated CI analysis: Graph Filtered Temporal (GraFT) dictionary learning. The graph creates a data-driven space where the correlation of dictionary coefficients over space during dictionary learning becomes a natural regularizer no matter what the scale of imaging. While we motivate and apply this method here to optical imaging data, the method itself is very broad and falls into the general class of graph-regularized dictionary learning (e.g., [36], [37]). We thus first present the algorithm in its most general form, followed by demonstrations both in simulated data, and in real recordings of somatic and dendritic data.

## II. Background

### A. Matrix factorization for CI data

The predominant form of automated calcium imaging analysis is regularized matrix factorization [23], [27]–[29]. Following the model of Equation (1), these methods aim to solve the cost function

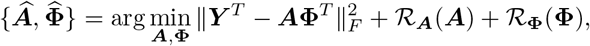

where here 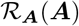 and 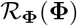 represent appropriate regularizations that can vary between methods, and often include terms such as the component norms, number of components, spatial and temporal sparsity, and spatial cohesion, as well as explicit modeling of the calcium dynamics [27]. As direct optimization is often difficult for problems of this size, alternating descent type algorithms are often employed, i.e., iteratively solving

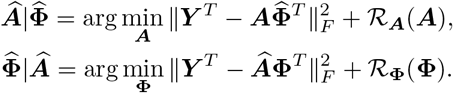

These methods can often be susceptible to noise and are, for example, particularly sensitive to initialization procedures and pre-processing (e.g., motion correction, baseline subtraction, variance normalization etc.). We instead propose to leverage a dictionary learning (DL) framework.

### B. Dictionary learning

DL is an unsupervised method aimed at finding optimal, parsimonious representations given exemplar data [38], [39]. Consider the model in Equation (2) and the equivalent Gaussian likelihood model of the data given the coefficients ***A***, i.e., 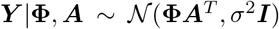, and assume a sparsity-inducing prior distribution *p*(***A***) over the coefficients ***A***. The invoked sparsity encourages learned representations that parsimoniously represent the data, i.e. each data-point can be reconstructed using only a small number of dictionary elements. DL in its purest form seeks to infer only the dictionary **Φ** via maximum likelihood, marginalizing over the coefficients

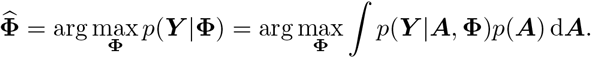

This marginal maximum likelihood optimization involves an often intractable integral. In particular, the prior *p*(***A***) is often chosen to be a *i.i.d.* Laplacian distribution 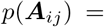 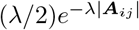. To bypass this difficulty, variational methods are often employed to iteratively solve this optimization [40]– [42]. For DL, this often takes the form of alternatively solving for the sparse coefficients of a subset Γ of the data, i.e.,

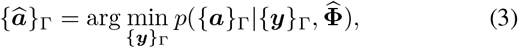

followed by the equivalent of a stochastic gradient descent over **Φ** using this subset of estimated coefficients

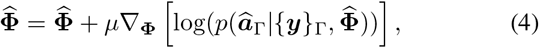

for some step size *μ*. These steps are often termed the “inference” and “learning” steps, respectively, and stem from an interpretation as inference of the maximum marginal likelihood via an expectation-maximization procedure [40]–[42]. Traditional sparse coding assumes this model with Dirac-delta posterior approximations and an exponential (Laplacian) prior resulting in the more commonly known inference/learning steps of

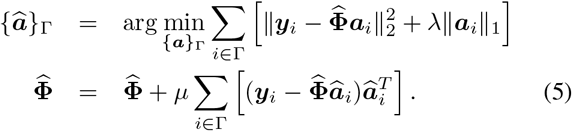

The capabilities of DL to learn generic features that efficiently represent data have found use more generally in applications beyond the original image processing applications [38], [43]. For example, in hyperspectral imagery (HSI) [44], [45], DL has been used to identify material spectra directly from image cubes in an unsupervised manner.

Previous work has also explored, to an extent, the application of DL to CI data [29], [46], [47]. Current applications focus on learning spatial dictionary elements [29], [46], [47]. These applications include spatial generative models based on convolutional sparse block coding [46], extensions of convolutional sparse coding to video data with non-uniform and temporally varying background components [47], and the DL of spatial components via iterative merging and clustering [29].

All these methods essentially isolate spatial dictionaries to lean neuron shapes, then using those shapes to find corresponding time courses. In our proposed method, Graph-Filtered Time-trace (GraFT) dictionary learning, we instead define the critical features as temporal features. To still leverage the important spatial information in a flexible way, we replace the spatial filter in spatial filtered sparse coding of [34], [48] with a graph-based diffusion filter.

### C. Sparsity-based Stochastic Filtering

The subselection of points Γ is often uniformly at random and thus DL does not inherently take into account relationships between exemplar data points. Recent models, however, provide new tools to modify the inference stage to take into account that some data-points may have similar decompositions [35], [48]–[50]. One such method, termed Reweighted *ℓ*_1_ Spatial Filtering (RWL1-SF) uses an auxiliary set of variables, ***λ***, to correlate the probability of coefficients in neighboring vectors being active [35], when applying DL to imaging / HSI data. Based on the reweighted-*ℓ*_1_ extension of basis pursuit denoising (BPDN) [51], [52], this method alternates between updating ***λ*** and ***a*** as

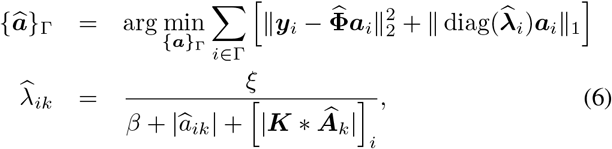

where 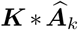 is the 2D convolution between a kernel ***K*** and the coefficient image for the *k^th^* dictionary coefficient 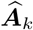. This convolution defines the local influence of the neighboring coefficients on the *i^th^* data vector’s decomposition weights. While previous work naturally fits into this deconvolution interpretation of a neighborhood [34], [35], in this work we expand on this idea and redefine what it means for two pixels to be “close”, using the formalism and tools of data-driven graph constructions and manifold learning.

### D. Graph Sparse Coding

A prevalent assumption in high-dimensional data analysis is the manifold assumption, according to which in many real applications the data, while being observed in a high-dimensional feature space, actually lies on or near a low-dimensional manifold embedded in the high-dimensional ambient space [53]–[55]. Under this assumption, when learning a new representation for data, two datapoints should have a similar representation if they are intrinsically similar, i.e. close to one another on the manifold. The manifold assumption has played an increasing role in dictionary learning via graph regularized sparse coding (GRSC) [56]–[60], where the underlying manifold is represented in the discrete setting as a graph on the data-points. Formally the graph is denoted 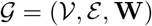, where 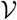 are the nodes in the graph, *ε* is the set of edges connecting the nodes, and **W** is the undirected weighted adjacency matrix for 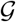, with **W***_ij_* ≥ 0 capturing the degree of similarity between nodes *i* and *j*. Let ***D*** be a diagonal matrix whose diagonal elements ***D**_ii_* = *d_i_* = Σ_*j*_ ***W**_ij_* are the nodes degrees of the graph. The symmetric graph-Laplacian is defined as ***L*** = ***D*** − *W*, and is used in the sparse coding inference step to “encourage” datapoints who are close in the graph to employ similar representations with respect to the dictionary. For example, the coefficient inference objective in [58] is given by

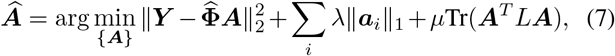

where 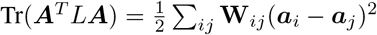.

In image processing applications, GRSC enables incorporating non-local structure [61] by considering similarities in the high-dimensional feature domain as opposed to only relying on local spatial information. In this paper, we propose a new sparse coding solution, which adapts re-weighted *ℓ*_1_ spatial filtering [35] to a graph-based filter. Instead of the symmetric graph Laplacian, we use the normalized random-walk kernel ***K*** = ***D***^−1^***W***. This kernel allows us to replace the spatial convolution kernel with diffusion along the data-driven graph constructed on the pixels. This kernel can be interpreted as a non-local averaging filter [62]. Next we describe the mathematical model and derive the algorithm that accomplishes these tasks.

## III. GraFT Dictionary Learning for Functional Imaging

As DL was initially presented in the image processing literature, early applications to CI focused on learning spatial features indicative of fluorescing cells and processes [29], [46], [47]. The benefit to learning a set of temporal dictionary vectors instead is that we directly model the main objects of interest to science: the neural activity traces. To infer these traces we consider the hierarchical model in Figure 1D, which describes the statistical relashonships between the time-traces **Φ**, data ***Y***, presence coefficients ***A***, weights ***λ***, and data graph ***K***. Mathematically we can define this model via the conditional probability distributions

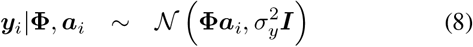

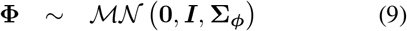

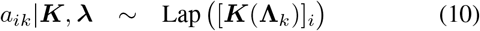

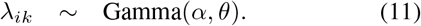

Starting at the top, Equation (8) places a isotropic mean-zero, variance 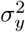 Gaussian likelihood on the fluorescence trace at each *i^th^* pixel given the temporal dictionary and coefficients. Next, in Equation (8), we place a mean-zero Matrix normal prior over the dictionary. The between-dictionary covariance matrix **Σ**_*ϕ*_ both penalizes unused time-traces (similar to [63]) and penalizes time-traces that are too similar. The prior over the coefficients (Eqn. (10)) takes the form of a sparsity-inducing Laplacian distribution where the parameter for the *k^th^* coefficient at pixel *i* is given by applying the *i* element of the function ***K***(**Λ**_*k*_), [***K***(**Λ**_*k*_)]_*i*_. This function applies the data-driven graph information encoded in the ***K***(·) to **Λ**_*k*_, the set of all weights for coefficient *k* across the entire image, to account for the correlations between data vectors at different pixels. The weights themselves, ***λ***, follow a conjugate Gamma hyper-prior with parameters *α* and *θ*, comparable to previous work [34], [35]. Our model represents a GraFT dictionary-based linear generative model where we can use a DL approach to learn the time-traces and graph-correlated sparse coefficients.

Inference under the above model can be complex and computationally intensive. We thus break down inference into three main stages: graph construction, coefficient inference and dictionary update. These stages are performed sequentially, as outlined in Algorithm 1 in an alternating minimization procedure. At a high level we are iterating over Equations (3) and (4). The main differences are a) we infer the coefficients for *all* data-points simultaneously, b) we replace the *i.i.d.* model with the graph-correlated model above, and c) we optimize the dictionaries completely at each iteration, rather than taking a single gradient step.

### A. Re-weighted *ℓ*_1_ Graph Filtering (RWL1-GF)

The first sub-step needed is to infer ***A*** under the model of Equations (8)–(11) given a fixed dictionary **Φ**. This involves expanding the idea of spatially filtered sparse inference (6) beyond the previous definition of neighborhood. Mathematically, we are moving from a convolutional kernel-based function 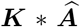 to the more general function 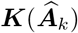 that defines the influence of the neighboring coefficients on the *i^th^* data vector’s decomposition weights for each dictionary index *k*. This optimization requires a definition of neighborhood, which previous convolution-based weights define using the spatial distance between pixel locations [34], [35]. In general applications, beyond imaging, it is difficult to define a universal pre-set sense of neighborhood, prompting the use of data-driven graph constructions. Under this formulation, 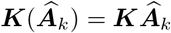 which is the result of multiplying the graph matrix ***K*** with a vectorized coefficient map / spatial map of the *k^th^* component. This can be interpreted as applying a single graph diffusion step to the coefficients.

There are multiple ways to construct a data-driven graph. Here we construct a graph on the pixels using *k*-nearest neighbors graph with the Euclidean distance between the time-traces ***y**_i_*. We calculate the weighted graph affinity matrix **W** using a Gaussian kernel for similarity, and set

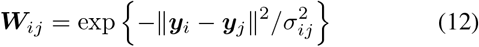

if ***y**_i_* is among the *k* nearest-neighbors of ***y**_i′_* or vice-versa. The bandwidth *σ_ij_* = *σ_i_σ_j_* is set to be the self-tuning bandwidth [64], which is a local adaptive bandwidth. Then the diffusion graph filter is calculated by normalizing the rows of ***W*** = ***K*** = ***D***^−1^***W***, where ***D*** is a is a diagonal matrix and ***D**_ii_* = Σ_*j*_***W**_i_*

Given the filter, we can solve the maximum likelihood inverse problem defined by Equations (8), (10) (11) for ***A*** via minimizing the negative log-likelihood

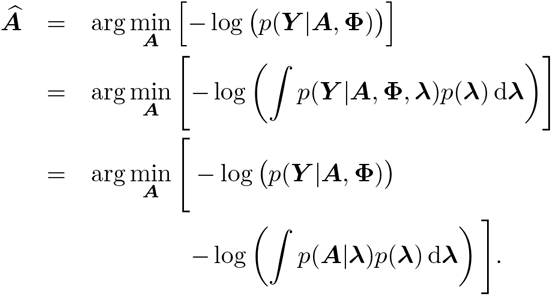

While exact integration of the above integral (marginalizing over ***λ***) can be very well approximated by the closed form

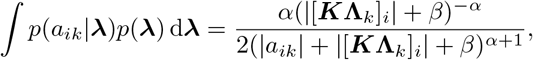

this prior results in a non-convex optimization negative log-likelihood cost function.

The sparse coding solution thus iteratively solves a weighted LASSO that can be interpreted as an approximation to a variational expectation-maximization (EM) optimization of the true ML problem [48], [52]. In this EM scheme, which we term Re-weighted *ℓ*_1_ Graph Filtering, the weights are updated at each algorithmic iteration based on smoothing with the graph-based kernel:

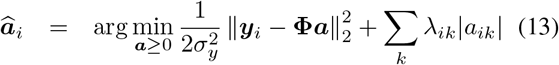

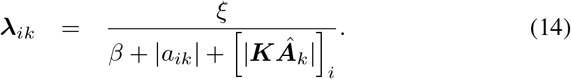

The weights *λ* in RWL1-GF incorporate spatial information into per-pixel solutions by sharing second-order statistics. Note that this spatial information is *non-local*, thus handling both components with compact spatial support such as cell-bodies, in addition to far ranging spatial components such as dendrites.

### B. Updating the temporal dictionary

The second step in the dictionary learning procedure is to update the time-traces themselves with respect to a learning rule. We use the above probabilistic model to specify an appropriate cost function. In particular, the matrix normal prior induces two regularizing terms to the cost. The first penalty involves the Frobenious norm over **Φ**, essentially penalizing excess activity. This term should, and we show in later experiments that it does, remove unneeded time-traces from the dictionary by setting them to zero. Thus the exact number of components need not be known *a prior* and can be modified autoLmatically. The second penalty, 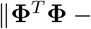 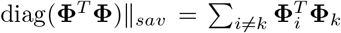 is a function of the intra-dictionary correlations^3^. This penalty ensures that time-traces are not learned with multiple times with minute differences stemming from subtle nonlinearities or noise. These terms come together in the cost function

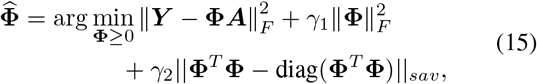

where the parameters *γ*_1_ and *γ*_2_ trade-off these penalties. These parameters have a direct link to the covariance matrix **Σ**_*ϕ*_ in Equation (9) when **Σ**_*ϕ*_ is the sum of the identity plus a rank-one matrix.

We note that in our formulation we are optimizing the entire dictionary completely at each iteration. This is because we are inferring all pixels mixing coefficients at each iteration in order to maximally utilize the complex graph connectivity. We thus lose some of the robustness imbued by a stochastic gradient descent procedure, and therefore modify Equation (15) to ensure stable convergence. Specifically, we include a continuation term that penalizes the change in the estimate 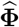 between updates. This term prevents too large a deviation in the dictionary between iterations, smoothing out the cost landscape and allowing for more robust solutions.

Mathematically, the update for the estimate at the *t^th^* iteration 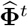 is given by

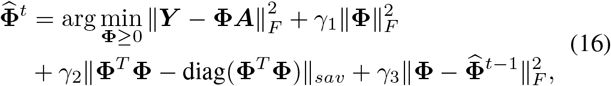

where *γ*_3_ is the parameter determining the rate of change in the dictionary over iterations.

### C. GraFT Dictionary Learning

The full algorithm incorporating all these elements — Data-driven graph correlations, re-weighted *ℓ*_1_ coefficient estimation, and robust, regularized time-trace learning — comes together in Algorithm 1. Specifically, we initialize with a graph constructed from the raw data temporal correlations and a randomly generated dictionary. The algorithm then iterates between using the new RWL1-GF algorithm to find approximate presence values, and the dictionary learning optimization of Equation (16). The final output is the learned dictionary 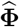.

#### Algorithm 1 GraFT DL

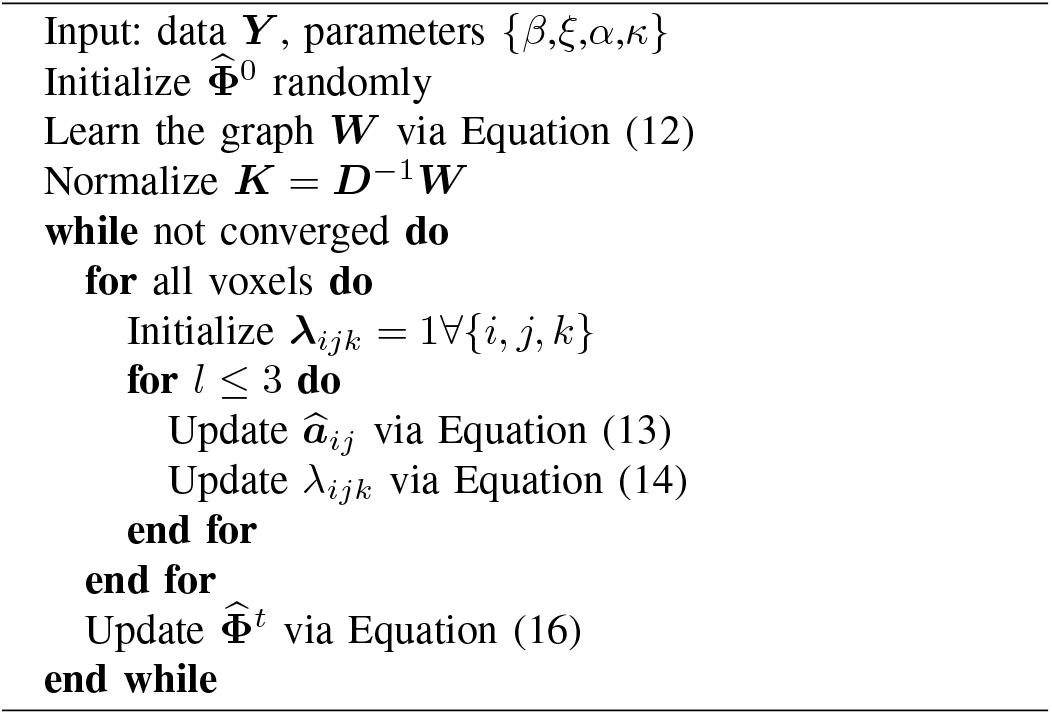

## IV. Results

To test GraFT we assess its performance both on simulated benchmark data, as well as apply GraFT to multiple datasets across all scales of optical imaging. Specifically, we assess the utility of GraFT for somatic, dendritic and widefield data.

### A. Implementation considerations

A number of practical considerations arise in our method, specifically initialization, selecting the number of neurons, setting the graph connectivity *W*, and parameter selection. First, we initialize **Φ** with random values, demonstrating a reduced sensitivity to initialization than other approaches [27]. Second, the number of dictionary components should be set to more than the expected number of neurons and background components. The sparsity and Frobenius norms serve to decay unused components (implicitly estimating the number of neurons), but cannot add new elements. To construct the graph pairwise affinity matrix *W*, we use a k-nearest neighbor construction. Each pixel is connected to *k* = 48 nearest neighbors and the affinity matrix is symmetrized. For parameter selection, we manually adjusted parameters using intuition build from prior work in RWL1-based algorithms [34], [35], [48]. The full set of parameters to set are *λ*_0_, *λ*_1_, *λ*_2_, *λ*_3_, *ξ*, and *β*. We find that for appropriately normalized data (normalized to the the median pixel value across the entire dataset), *λ*_2_ = *λ*_3_ = 0.1 provide accurate results across all datasets. Similarly we find that *λ*_1_ = 0.2 appropriately penalizes extra components. For the RWL1 parameters, we set *ξ* = 2 and *β* = 0.01 for all experiments. This choice sets a maximum cap of 200 for the per-element sparsity parameter modulator, and enables the weights to meaningfully vary as ≈ *a*^−1^ for small values ≈ 1. *λ*_0_ is the main parameter we find necessary to vary and in our experiments we manually tune this parameter in the range 0.001 < *λ*_0_ < 0.1.

Operating on the entire dataset simultaneously can often be time-consuming. Thus, for somatic datasets we follow similar procedures in other pipelines whereby the full field-of-view is partitioned into smaller, overlapping spatial partitions [27], of size ~ 50 × 50 pixels with 5 pixels overlap between partitions. The number of dictionary atoms in each partition is set to be on the order of 5-10. Each partition can be analyzed in parallel, and the results are merged together to ensure that components present across partitions are appropriately combined. Merging is accomplished by weighted averaging of highly correlated dictionary components (> 0.85), and then recalculating the spatial maps over all pixels.

A final consideration is the pre-processing that most algorithms use to denoise data before analysis. Often spatial and temporal low-pass filtering is performed. To maintain high-frequency temporal content, we instead denoise each pixel’s time-trace independent of all other pixels with a simple wavelet-based Block James-Stein (BJS) [65] denoising in a 2-level, ‘sym4’ wavelet decomposition. With these aspects in mind, we implemented our code in MATLAB^4^.

### B. NAOMi Simulated two-photon calcium imaging

To test our framework, we use recent advances in highly-detailed biophysical simulations that produce realistic two-photon imaging data based on computational models of anatomy and optics, i.e., the NAOMi simulator [31]. As the anatomy and time-traces are all simulated, the full spatio-temporal ground truth is known, including voxel/pixel occupancy of neurons (the spatial profiles of every fluorescing component), and the changes in fluorescence over time (each component’s time-trace). Rather than being “drawn from the generative model”, this data is generated by sampling 3D volumes of neural tissue and calcium activity, and then simulating the optical propagation and sampling that generates the CI data.

We applied GraFT to a 500 *μ*m × 500 *μ*m simulated field-of-view with 20,000 simulated frames. The simulation framerate was 30 Hz, and the spatial resolution was 1 *μ*m/pixel. The resulting found components were compared to results using the three most popular algorithms for ROI extraction, CNMF [27], Suite2p [26], and PCA/ICA [23]. As a further point of comparison, we compute the profile-assisted least-squares (PALS) time-traces based on oracle knowledge of the spatial maps. PALS uses the ground-truth spatial maps given by the NAOMi simulation, ***A***^†^ and computes the least-squares time traces as

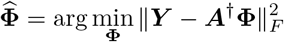

While PALS is not ground truth (the ground truth traces are independently provided by the NAOMi simulation), the traces PALS provides are an approximate baseline of per-neuron SNR and the general visibility of neurons in the dataset sans any regularization. We note, however, that they are not oracle traces, as PALS leverages no regularization or significant pre-processing (e.g., the denoising present in all other algorithms). When comparing the found components to the ground-truth traces, we find that GraFT identifies 434 unique neural components (i.e., including somatic and dendritic components), 37.76% more true positives than the next best performing algorithm, CNMF, that found 303 unique neurons, and even 19 (4.58%) more than the PALS time-traces.

### C. Two-photon population calcium imaging

In addition to assessing GraFT on simulated data, we furthermore test GraFT on data from the NeuroFinder [66] dataset. NeuroFinder data includes two-photon calcium imaging at the somatic scale, along with manually annotated spatial maps for neurons. We note that while these labeled spatial maps are provided, they are limited in their use as ground truth. Some neurons are visible but do not fire and are thus functionally unidentifiable, while others active neurons are not labeled. In addition, other clearly fluorescing components might not be labeled as they may represent dendritic (apical or otherwise) components. We ran GraFT on a 445 *μ*m × 445 *μ*m field of view in area vS1 of mouse visual cortex, recorded at 8 Hz. For comparison we ran Suite2p, optimizing parameters of both algorithms manually.

In total, GraFT and Suite2p identified 150 and 103 components respectively. GraFT and Suite2p found 46 and 48 out of 56 labeled components respectively (Fig. 3A-C). GraFT, however, did identify a number of extra neuronal components, including somas, dendritic segments and apical dendrites. Further inspection revealed that movie frames during active bursts in the learned dictionary time-traces matched the spatial profiles, verifying that these components do represent real fluorescing components in the data (Fig. 3D). Identifying these additional components individually removes their localized activity from the soma-adjacent areas, improving neuropil estimation and subtraction.

**Fig. 2.**
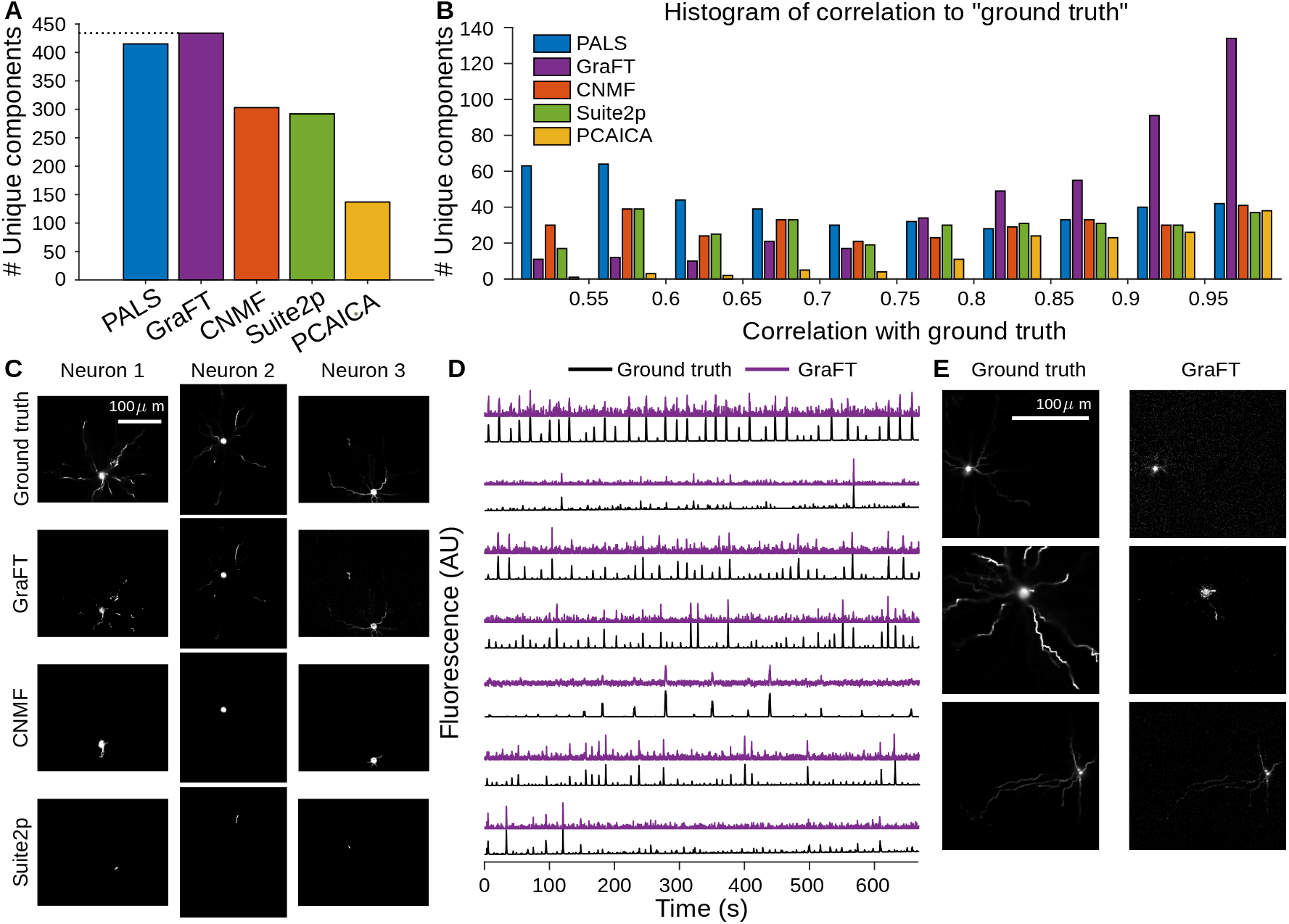
Assessment of GraFT on anatomically-based calcium imaging simulations and comparison to current methods. A: The number of unique neurons in the dataset found with each method including PALS, which represents the SNR of each time-trace. B: Histogram of temporal correlations between the time-traces of found neurons and the ground truth traces demonstrates that the GraFT dictionary better matches the ground-truth, compared to other methods. C: GraFT finds more complete spatial profiles for neurons also identified by other methods. In particular dendrites are better identified. D: Time-traces for neurons not found via other methods still correlate well with ground truth traces, but have low SNR. E: Neurons found only with GraFT tend to have less localized spatial profiles.

**Fig. 3.**
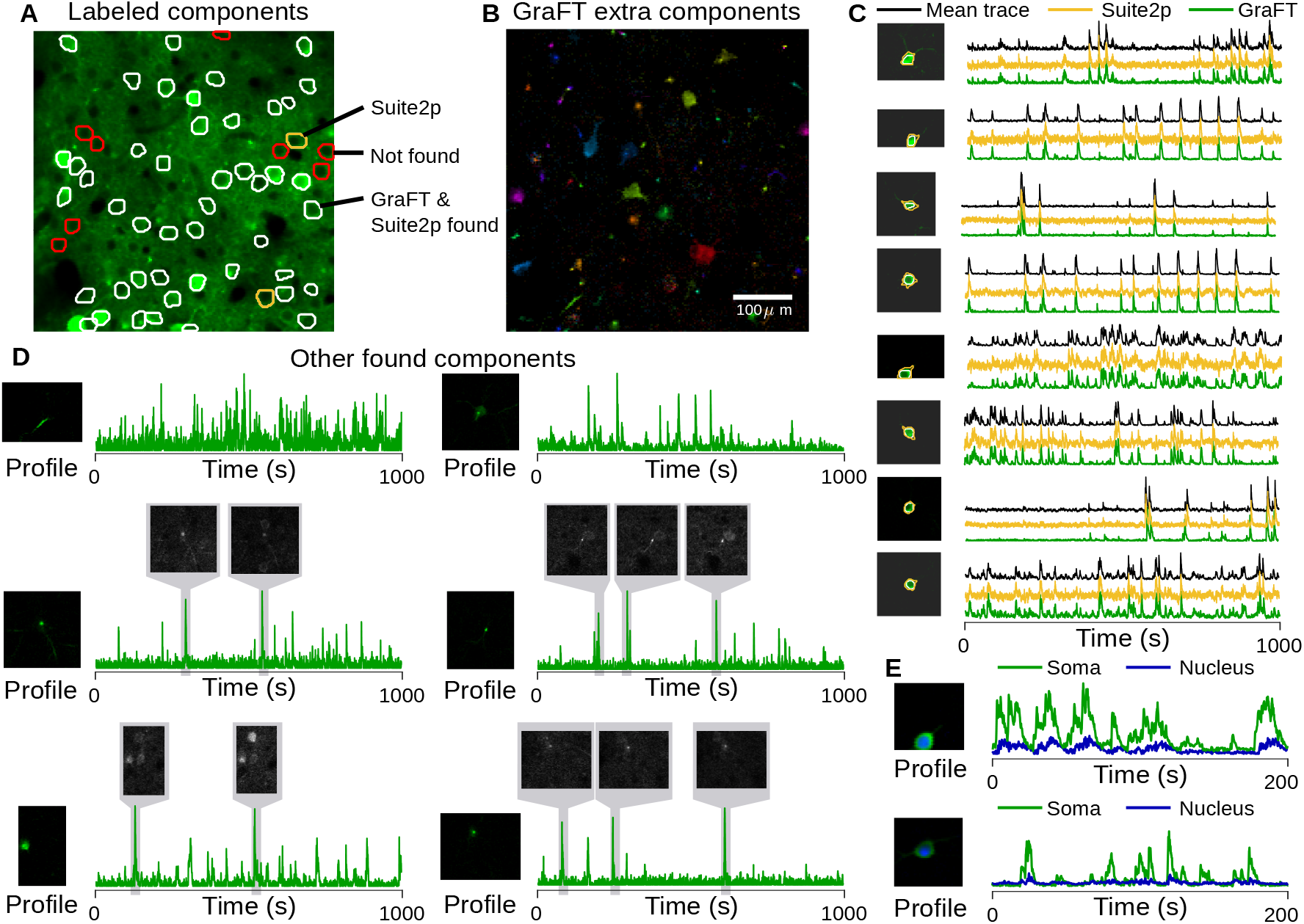
Assessment of GraFT on somatic imaging from the NeuroFinder dataset. A: Imaged field-of-view with all manually identified cells circled and overlaid on the mean projection of the data (green). White circles indicate profiles that both GraFT and Suite2p identified, red indicates profiles that no algorithm identified (i.e., due to inactivity), and yellow indicates cells that Suite2p found that GraFT did not. Note that even in the mean projection there are obvious bright cells and dendritic components not labeled as ground-truth. B: Image of all non-background (well-isolated) profiles found by GraFT that were not in the manually labeled set of data. Included are many apical dendrites, several somatic components and other small components. C: Examples comparing the time-traces found via GraFT and Suite2p with the average trace over the manually identified spatial profile. We note that the GraFT time-traces tend to have less noise, despite not explicitly regularizing for smoothness. Suite2p traces tend to exhibit negative “dips” indicating poor neuropil correction. D: Additional components beyond the manually identified profiles exhibited smaller, dendritic structure, even when these are in proximity to brighter somatic components (right column, middle). Examination of local averages of the calcium imaging movie demonstrate that these components do, in fact, exist in the data and were simply not identified manually. E: The sensitivity of GraFT enables for the cytoplasmic and nuclear portions of individual neurons to be identified. Two examples show the cytoplasmic (green) and nuclear (blue) components. As expected, the nuclear signal is lower amplitude and has slower dynamics.

Finally, we noted that some somatic components appeared to be found in duplicate. We explored these cases and discovered that GraFT was, in fact, separating the cytoplasmic and nuclear portions of these cells (Fig. 3E). When plotted together, the time-traces for each pair follow closely together, with a muted and delayed response for the nuclear component relative to the cytoplasmic. Thus, GraFT is able to partition even very similar activity patterns, revealing potentially interesting differences within individual cells.

### D. Sparsely labeled dendritic imaging

One of the principle benefits of GraFT is to enable the extraction of non-compact components in the data. While somatic-scale imaging contains some dendritic components, dendritic-specific imaging at high zoom levels drastically changes the spatial statistics of the data. We thus next apply GraFT to dendritic imaging of both sparsely and densely labeled tissue. First we use the sparsely labeled data to assess the accuracy of GraFT via comparisons to anatomical measurements.

We imaged dendritic calcium signals in an awake sparsely-labeled mouse using a Bruker 2P-Plus microscope with an dual-beam Insight X3 laser (Spectra Physics), equipped with an 8 kHz bidirectional resonant galvo scanner and a Nikon 16X CFI Plan Fluorite objective (NA 0.8). Fluorescence was split by a 565LP dichroic and filtered with 525/70 and 595/50 bandpass filters before collection on two GaAsP photo-multiplier tubes (Hamamatsu H10770PB-40 and H11706-40, respectively). We imaged a square region 375 *μ*m per side, 768 × 768 pixels. Frames were acquired at 20 Hz and 13-bit resolution. Illumination was centered at a wavelength of 940 nm, and laser power exiting the objective was in the range of 30-40 mW. PMT gains were set to minimize saturated pixels during calcium transients.

We supplemented the functional imaging with anatomical imaging to provide ground truth for assessing the GraFT decompositions. Immediately following calcium imaging experiments, we anaesthetized the mouse with 1-2% isoflurane and placed a heating pad underneath the mouse. We then recorded large field-of-view volumetric z-stacks (750 *μ*m square, 1536 × 1536 pixels, 5 micron axial spacing) of mRuby2 fluorescence by illuminating at 1045nm using the galvo scanners at a 10 microsecond dwell time. This z-stack was manually traced, to recover a depth tracing of the dendrites of each of the individual neurons in the FOV, which serves as spatial ground-truth for assessment. The depth tracing reveals 6 neurons, 3 of which were active in the functional imaging data and identified by GraFT.

As a preprocessing step, we apply a structural mask to the imaging video which yields 91,737 pixels in the FOV. The mask was constructed by thresholding a mean projection of the mRuby2 fluorescence channel from the functional imaging session. For sparsely labeled data, GraFT identified 10 distinct components in the data (Fig. 4A,B). Seven of these components are well localized with distinct firing patterns, and three are baseline/background components. Comparisons to the anatomical image stack show that all found components match well to at least one of the identified neurons in the volume (Fig. 4C). Interestingly, some components appeared to capture different spatial portions of the same dendritic branch. We further examined this effect and noted that these components are capturing the same anatomical object but at depth difference of approximately 10-15 *μ*m (Fig. 4D). Thus GraFT is identifying the same anatomical component, but modulated due to the slow axial drift in the plane of imaging.

**Fig. 4.**
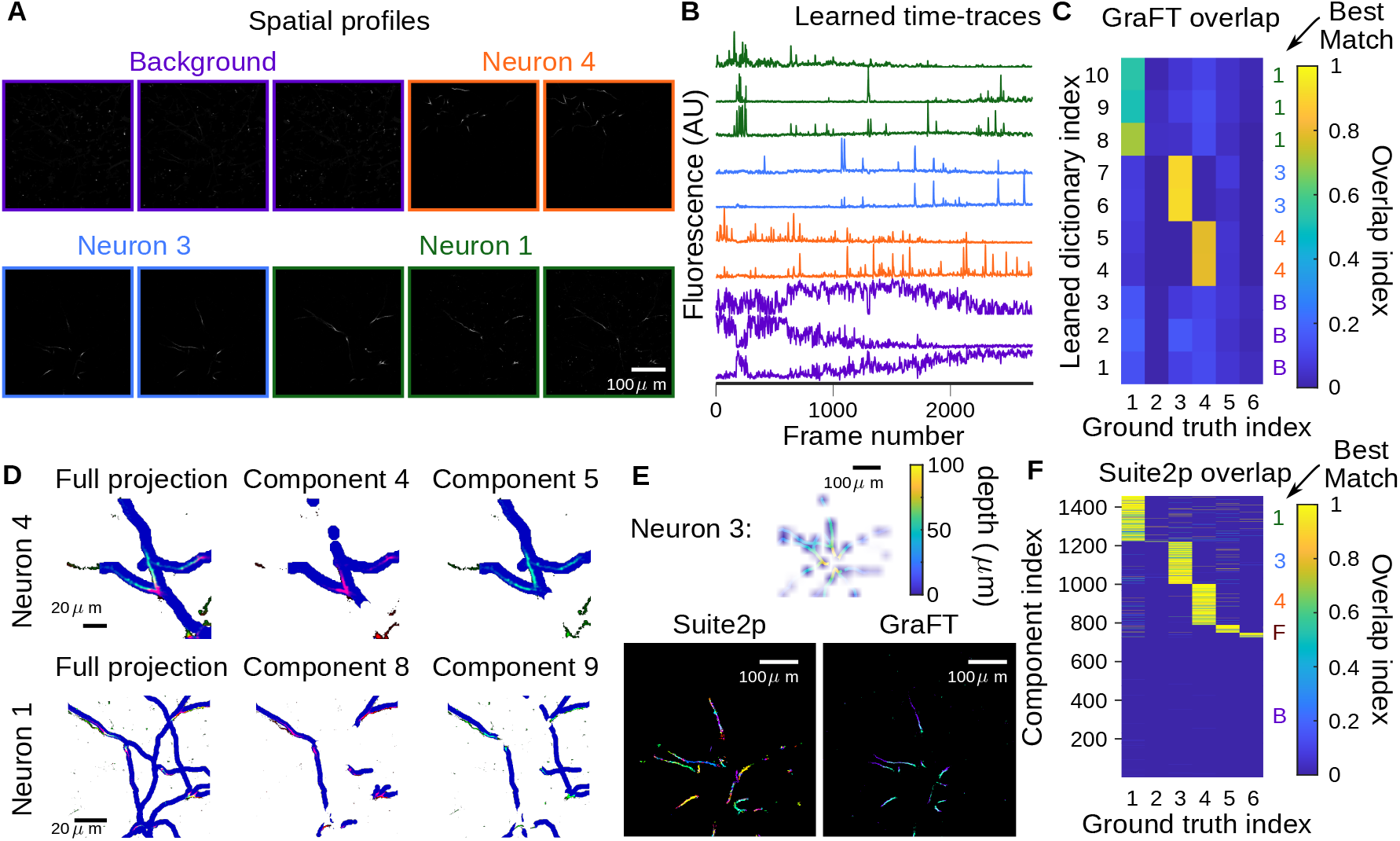
Assessment of GraFT on sparse dendrite data. A-B: 10 temporal dictionary elements along with the 10 corresponding spatial maps recovered after running GraFT with 10 components. Dendritic components can be seen extending throughout the entire image, as is expected. C: Correlation matrices show 2 types of components were found: those serving as background components with approximately equal overlap across all dendrites, in contrast to those with very high overlap with only one anatomically traced neural dendrite structure. Interestingly, two-three components were found for each of the ground truth components. Note that two neurons that were not active at all and were not identified in any of the learned components. D: Analysis of the two components found per ground truth structure reveals that slightly different portions of the neuron (dilated for effect in blue) are captured by each of the two components matched (colored in red and green channels). Closer inspection reveals that each component corresponds to a slightly different axial portion of the neuron, indicating that the difference is due to axial motion. This effect is shown here for two of the three neurons in the volume, with the full ground truth projected onto one image on the right column, and the middle and right plots showing the ground truth at slices shifted by 2 *μ*m. E: Comparable methods, for example Suite2p run in “dendrite” mode tends to use many more components to describe a single dendrite. For example, for Neuron 3, approximately 200 ROIs (different colors in the lower-left plot) comprise this single neuron. GraFT picks up the same neuron with two components: each corresponding to a different depth during the axial motion discovered in panel D. F: The effect of Suite2p using many components to cover a single dendrite was observed for all neural components, as well as the background component. Overall over 1400 components were extracted (with 200 ROIs per active neuron) versus the 10 spatial maps GraFT extracted.

Since GraFT does not impose that the spatial maps need to be contiguous or spatially connected, we are able to extract disconnected segments of the same dendrite within a single spatial map. In comparison, Suite2p extracts 1,457 ROIs (note that we removed all ROIs Suite2p found that did not overlap with the structural mask used for GraFT). Of the extracted ROIs approximately 650 correspond to the 3 active neurons in the FOV, with about 200 ROIs matching each neuron (Fig. 4F-F). GraFT thus easily enables studying the dynamics of the dendrite as a whole, whereas Suite2p requires post-processing by which ROIs need to be clustered together based on temporal correlations in order to study the dendrite as whole.

### E. Dense dendritic imaging

While sparse imagine provides the possibility of comparisons to anatomical imaging and tracing, we are also interested in the ability of GraFT to disentangle components in images of tissue with a higher density of labeled dendrites. We thus run GraFT on a dataset of densely labeled dendrites using the same imaging parameters as the sparse dataset. Overall GraFT extracted 60 individual components from the dataset, including many with spatial maps stretching across the entire field-of-view (Fig. 5A). In comparison, Suite2p extracted approximately 800 ROIs (results not shown). The time-traces for the GraFT components are more uniformly firing over the span of the imaging session, indicating a lack of axial drift that was noticed in the sparse imaging data (Fig. 5B). Furthermore, these components are significantly diverse in their time-traces, as can be seen in the correlation matrix between time traces (Fig. 6C). One interesting finding in this decomposition is the presence of multiple time-traces identified in the same dendritic branch (Fig. 6D-E). Specifically, for one pair of components we find that one of the components is much more localized around one small branch, specifically about one spine (Profile 2 in Fig. 5D). The time-traces for these two components, while correlated, do have different activity levels as can be verified by tracking the components in the raw fluorescence video (Fig. 5E).

**Fig. 5.**
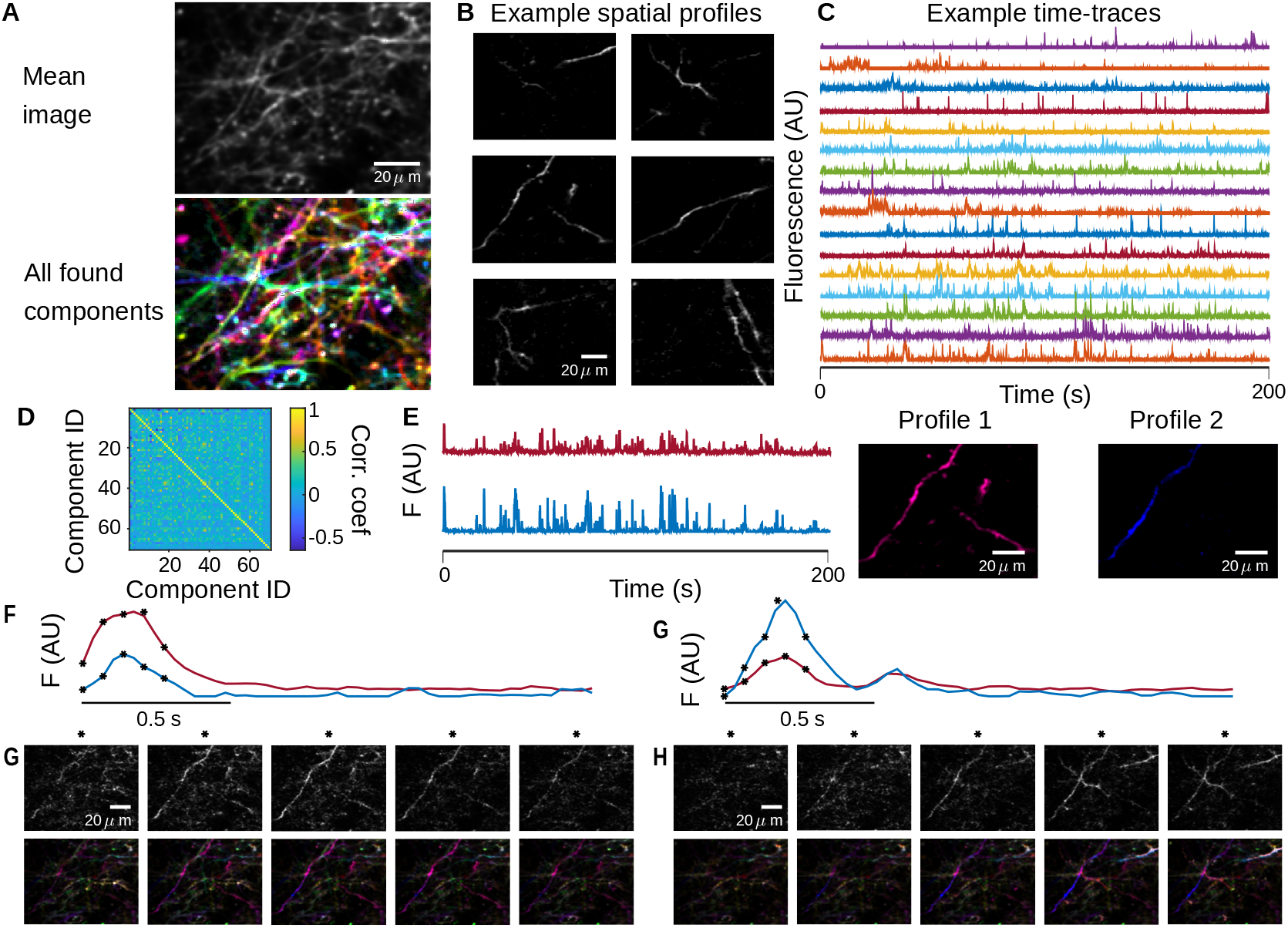
Assessment of GraFT on dense dendrite data. A: In the dense dendritic imaging, many fluorescing components are highly overlapping, as can be seen in the movie’s mean image (top). GraFT identifies 70 components in this dataset (bottom image; color coded with each found component as a different color), which span the entire field of view. B: Examples subset of spatial maps recovered after running GraFT with 60 components. C: Example time-traces demonstrate a diversity of activity patters that overlap in time. D: The correlation matrix, combined with the diversity of time-traces in B, indicate that the decomposition is capturing sufficiently different time-traces. E: The two highest correlated time-traces are shown, along with the corresponding spatial maps. At first glance it appears that these two spatial maps overlap significantly, perhaps both representing pieces of one true dendritic component. F-G: Closer inspection of the spatial profiles reveals that profile 2 actually represents a different process in the dendrite centered around a spine in the lower-left-hand corner of the image. Two example bursts of activity (E and F) demonstrate that in fact these components do show up in different quantities at different times. Frames from the starred time-points in E and F are shown in G and H respectively, with the raw data frame in the top row and the reconstruction in the bottom row. In G Profile 1 appears brighter, as implied by E and in H Profile 2 appears brighter, as implied by F. Thus these profiles are truly independent components and not an artifact of the method.

**Fig. 6.**
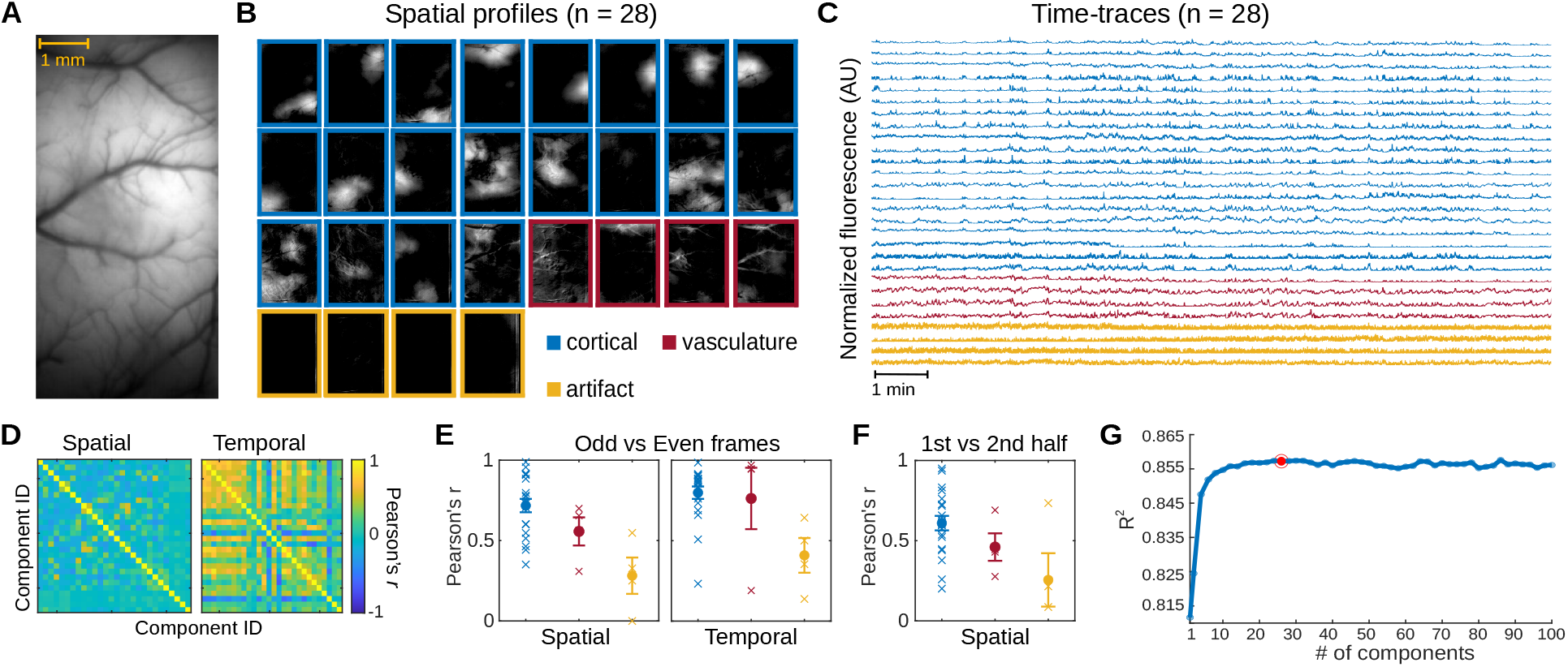
Application of GraFT to widefield data. A. The imaged cross-cortical field-of-view. B. GraFT spatial maps when run with *n* = 28 components. Three main classes of components can be identified: broad cortical activity (blue), vasculature-based fluorescence (red), and imaging artifacts (yellow). C. Time-traces for the *n* = 28 case, color coded by the class of the component. D. GraFT demonstrates inference consistency, as demonstrated by comparing the learned time-traces and spatial maps for GraFT run only on the odd and even frames separately. The Pearson’s correlation between best-matched spatial and temporal maps are very high for all maps and dictionary elements for the case of *n* = 28. E. Breakdown of the diagonal elements of the correlation matrix in D by class (cortical, vascular, and artifactual). The learned spatial maps and time-traces for cortical maps was the most consistent, with the highest correlation values. F. A similar analysis of spatial maps learned separately across the first half and the second half of the imaging session for *n* = 28 demonstrates that consistent cortical areas are observed during the same behavior even over different epochs. G. Variance explained as a function of the number of dictionary elements shows a plateau at approximately 0.855 with ≈ 20 components.

### F. One-photon widefield imaging

Similar to dendritic imaging, widefield imaging captures activity patters that can stretch across the entire field-of-view. Widefield imaging systems can be designed to capture dynamics across large areas of the surface of cortex at a resolution coarser than single neurons. Thus widefield captures global activity patterns across the brain’s surface, similar to very high density EEG, giving us a second imaging modality to test GraFT’s ability to capture complex neural activity patterns. We applied GraFT to a 20-minute long video of calcium dynamics within an 8 mm × 4 mm FOV of the dorsal cortex of a GCaMP6f expressing transgenic rat. The recording was performed using a head-mounted widefield microscope [14] as the rat freely moved around its homecage (Fig. 6A). With 28 components, GraFT extracted a diverse set of dynamics from the widefield data (Fig. 6B-C). Most of the learned components featured spatial profiles that are either localized or widely distributed across the cortical surface (*n* = 20/28). Pixels belonging to observable vasculature were assigned low weights among these spatial profiles. In contrast, four other components showed spatial profiles with vasculature structures. The remaining components have spatial profiles with vertical stripes in the peripherals of the FOV suggesting they represent artifacts related to the imaging procedure. The 28 components exhibit unique spatial profiles but have time-traces that correlate to varying degrees (Fig. 6D). To test the reliability of GraFT in extracting relevant dynamics in wide-field calcium imaging data, we trained GraFT on odd and even frames of the widefield video separately (Fig. 6E). The learned components exhibit spatial profiles that overlapped between the odd and even frames (average Pearson’s *r* = 0.6324), indicating that GraFT can reliably recover dynamics in the data only from a portion of the total frames. We then trained GraFT on the first and second halves of the widefield video (Fig. 6F). The average coefficient along the diagonal of the spatial correlation matrix between the first and second half of the video is equal to 0.5376, suggesting that GraFT was able to extract sufficiently stable components in the video. We further assessed the performance of GraFT in extracting components in the widefield data, by measuring the R-squared between the original video and reconstructions based on the learned components. GraFT was able to capture a significant proportion of variance in the original data with only one component (*R*^2^ = 0.812) and achieved maximum performance with 44 components (*R*^2^ = 0.858). While the performance of GraFT increases with additional components, the change in performance seems to greatly decrease with ≈ 20 components (Fig. 6G). Together, these results indicated that GraFT reliably captures the spatial and temporal calcium dynamics recorded via widefield imaging in freely moving rodents.

## V. Conclusions

We propose a new algorithmic framework for dictionary learning based on data-driven graphs. These graphs permit the learning process to explicitly take advantage of correlated occurrences of features in the data. Specifically, we combined ideas from spatially correlated re-weighted-*ℓ*_1_ filtering with random walk diffusion on graphs to create a re-weighted-*ℓ*_1_ graph-filtering (RWL1-GF) inference algorithm. Learning of the linear generative model under the RWL1-GF model was performed via a variationally-motivated dictionary learning algorithm and was able to uncover the fundamental time-courses in spatio-temporal data with complex spatial correlations.

This powerful framework could have many potential uses, however we design and demonstrate this algorithm on the important and rapidly developing application of functional fluorescence microscopy. We applied GraFT to learn the time-traces from these complex and high dimensional data that are vital to understanding the neural function. In particular we showed benefits both in standard somatic imaging, as well as the more complex dendritic and widefield imaging.

For somatic imaging we demonstrated via experiments on biophysical simulations the ability to identify many more true components in a given dataset than competing methods. These gains primarily come from components with highly non-somatic statistics. Despite the flexibility to capture such components, GraFT can still capture somatic signals, as demonstrated on both simulation and annotated Neurofinder data. In both cases the time-traces obtained by GraFT were less noisy despite having no temporal regularization or explicit modeling of the calcium dynamics (e.g., an autoregressive model as in [67]). The lack of explicit temporal modeling further endowed GraFT with a sensitivity to different signal time-scales. Specifically, in real data GraFT was able to identify and separate the cytoplasmic and nuclear signals within the same cell. These signals essentially have the same spike-train driving both signals, albeit with different temporal signal dynamics.

In dendritic imaging we validated the ability of GraFT to recover fluorescing dendrites via comparisons against manual anatomical tracing. Despite the highly non-localized nature of the components, GraFT found multiple true components. Current methods, i.e., Suite2p in “dendrite mode”, broke up each dendrite into many components (> 200/dendrite). Moreover, GraFT also was able to demix much denser labeled dendritic imaging data, including finding multiple modes of activation within the same neural structure. Similarly, in wide-field imaging GraFT was able to identify highly non-localized activity patterns in the data, including both hemodynamic and calcium related signals. Moreover, GraFT was able to isolate artifacts of the imaging procedure, effectively removing their effects from the true signal components.

With regards to parameter setting we observed that the same basic parameters, with the exception of the sparsity parameter *λ*, produced good results across all imaging scales.

This observation reinforces the ability for the graph learning step to incorporate the natural spatial correlations into the same algorithmic core. With one parameter left, manual adjustments or simple grid searches were sufficient to obtain good performance. Further improvements can be obtained by leveraging, for example, BayesOpt [68], in optimizing parameters.

One aspect we note, that is not solved in any algorithm to date, is the effect of axial motion artifacts on the identified components. In the sparse imaging session we are able to identify manually when the resulting components represent the same neurons at different depths due to drift, however a full solution would require significant additional post-processing to the model that include 3D spatial information.

GraFT, as with other segmentation algorithms, identifies multiple types of components. For example, in the Neurofinder data GraFT identified somas, nuclei, dendritic segments and rising apical dendrites. Currently, sub-selecting based on component type is a per-analysis choice, however future work (e.g., the post-processing classifier in CaImAn [67]) can provide additional algorithms that automatically classify the output of GraFT and other algorithms. As fluorescence microscopy continues to expand, for example in new volumetric imaging techniques across scales [12], [69]–[74], we expect GraFT to become even more critical to the important fist step of extracting single component activities.

Finally, our method is broader than fluorescence microscopy data and is applicable to other imaging modalities, e.g., hyper-spectral imaging, where spatial structure can be informative, but features can interact in complex ways. In such cases, the dictionary is in the feature space, and not necessarily temporal, e.g., spectral bands in hyper-spectral imaging. Our framework also falls under the general class of graph-based dictionary learning in graph signal processing, an emerging area focused on the analysis of high-dimensional signals that lie on a graph, or for which a graph can be learned based on correlated features. We leave the extension of our framework to additional domains for future work.

## Acknowledgments

The authors thank Samuel Schickler for running the comparison experiment of Suite2p on sparse dendritic imaging.

## Footnotes

1 We prefer to use the terminology “spatial profiles” over the alternate term “Regions of Interest” (ROIs), as we believe it more accurately captures the physical nature of these shapes as 2D projections of 3D anatomical shape.

2 While the noise in CI is actually not Gaussian, at higher photon counts the noise is more Gaussian-like, and we, as many others, have found that Gaussian noise assumptions a practical simplification.

3 In this definition we use the *sav* or *sum-absolute-value* which is the direct analog of the *ℓ*_1_ norm for vectors, i.e., ‖***A***‖_*sav*_ = Σ*_ij_*|***A**_ij_*|. This is necessary as the ‖***A***_1_‖ has an alterLnate technical definition as the maximum *ℓ*_1_ norm of all the columns max_j_ Σ_i_| ***A**_ij_*|

4 Code implementation is available at https://github.com/adamshch/GraFT-analysis

